# Effect of Suppression of Rotational Joint Instability on Cartilage and Meniscus Degeneration in Mouse Osteoarthritis Model

**DOI:** 10.1101/2021.05.27.445937

**Authors:** Kohei Arakawa, Kei Takahata, Yuichiro Oka, Kaichi Ozone, Kzuma Morosawa, Saaya Enomoto, Kenji Murata, Naohiko Kanemura, Takanori Kokubun

**Author notes:** Corresponding author: Takanori Kokubun, PhD, Department of Physical Therapy, Health and Social Services, Saitama Prefectural University Saitama, Japan. Authors’ email addresses: KA,; KT,; YO,; KO,; K Morosawa,; SE,; K Murata,; NK,; T. Kokubun.

## Abstract

**Objective:** Joint instability and meniscal dysfunction contribute to the onset and progression of knee osteoarthritis (OA). In the destabilization of the medial meniscus (DMM) model, secondary OA occurs due to the rotational instability and increases compressive stress resulting from the meniscal dysfunction. We created a new controlled abnormal tibial rotation (CATR) model that reduces the rotational instability that occurs in the DMM model. So, we aimed to investigate whether rotational instability affects articular cartilage degeneration using the DMM and CATR models, as confirmed using histology and immunohistochemistry.

**Design:** Twelve-week-old male mice were randomized into 3 groups: DMM group, CATR group, and INTACT group (right knee of the DMM group). After 8 and 12 weeks, we performed the tibial rotational test, safranin-O/fast green staining, and immunohistochemical staining for TNF-α and MMP-13.

**Results:** The rotational instability in the DMM group was significantly higher than that of the other groups. And articular cartilage degeneration was higher in the DMM group than in the other groups. However, meniscal degeneration was observed in both DMM and CATR groups. The TNF-α and MMP-13 positive cell rates in the articular cartilage of the CATR group were lower than those in the DMM group.

**Conclusions:** We found that the articular cartilage degeneration was effectively suppressed by controlling the rotational instability caused by meniscal dysfunction. These findings suggest that suppression of rotational instability in the knee joint is an effective therapeutic measure for preventing OA progression.

## INTRODUCTION

Osteoarthritis (OA) is a joint disease associated with cartilage degeneration, synovitis, subchondral bone degeneration, and osteophyte formation,^1^ and its progression can cause significant pain and disability.^2,3^ OA affects more than 150 million people worldwide, and knee OA accounts for most cases. ^2,3^ OA is a multifactorial disease wherein aging, obesity, and trauma have been shown to play roles in its development.^4–7^ Among the known factors for OA, the increase in mechanical stress plays a crucial role in the development and progression of knee OA. Joint instability is known to result in abnormal mechanical stress that may cause knee OA.^8^ Joint instability causes synovitis and cartilage degeneration by increasing the expression of the inflammatory cytokine, tumor necrosis factor-α (TNF-α), and the cartilage catabolic factor, matrix metalloproteinase (MMP).^9–12^

The ligaments, muscles, and meniscus provide stability to the knee joint. In particular, the meniscus enhances bone compatibility and reduces the compressive stress in the knee joint.^13^ In addition, the meniscus controls the movement of the femoral condyle on the tibial plateau and is essential for flexion, extension, and rotation of the knee joint. Tibial rotation during flexion and extension occurs in a normal knee. However, it has been reported that abnormal tibial rotation occurs in patients with knee OA.^14–16^ They also experience meniscal degeneration,^17–19^ which may contribute to rotational instability. However, there are no studies that explain the mechanism of rotational movement in cartilage degeneration with meniscal dysfunction.

The destabilization of the medial meniscus (DMM) model is the most used experimental rodent model for knee OA. In the DMM model, secondary OA occurs due to the abnormal mechanical stress resulting from the destabilization and degeneration of the medial meniscus.^20,21^ This meniscal dysfunction causes joint instability and increases compressive stress. We hypothesize that the DMM model could reproduce the rotational instability and meniscal degeneration observed in patients with knee OA. We also created a new controlled abnormal tibial rotation (CATR) model that reduces the rotational instability that occurs in the DMM model. In this model, we used a nylon suture to control the abnormal rotation from outside the joint capsule based on our previous model.^22^ Thus, we aimed to investigate whether rotational instability affects articular cartilage degeneration using the DMM and CATR models, as confirmed using histology and immunohistochemistry.

## MATERIALS AND METHODS

### Animals and Experimental Design

This study was approved by the Animal Research Committee of Saitama Prefectural University (approval number: 29-13), and the animals were handled in accordance with the relevant legislation and institutional guidelines for humane animal treatment. The experimental design is illustrated in **Figure 1A**. 20 twelve-week-old male mice were procured for the study (Institute for Cancer Research), and a total of 30 knees were assessed in subsequent experiments. The mice were randomized into 3 groups: DMM group (DMM, n = 10), CATR group (CATR, n = 10), and no surgery group (INTACT, n = 10; right knee of the DMM group). All mice were housed in plastic cages under a standard 12 h light/dark cycle. Mice were permitted unrestricted movement within the cage and had free access to food and water.

**Figure 1.**
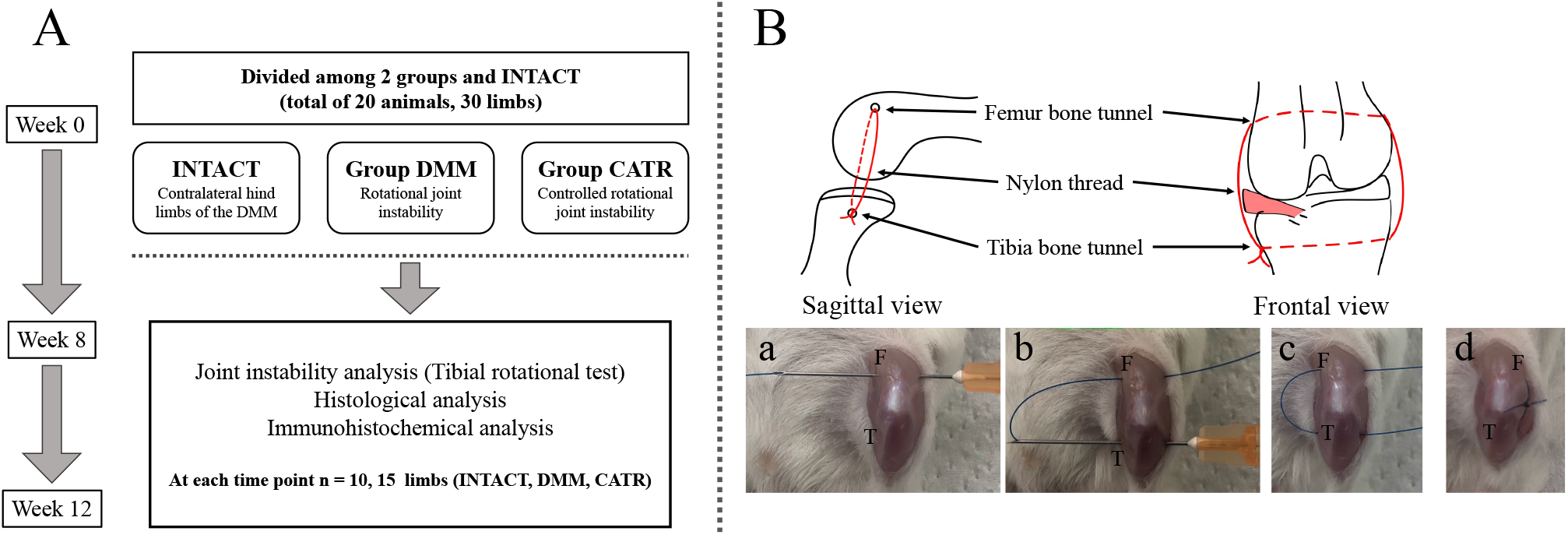
(A) Experimental design. We performed tibial rotational test, histological analysis and immunohistochemical analysis at 8 and 12 weeks. These analysis involved the DMM group, CATR group, and INTACT group (for each group, n = 5). (B) Surgery for the CATR model. A bone tunnel were created in the distal femur and proximal tibia using a 25-gauge needle (a,b) and insertion of 4-0 nylon threads threaded through the bone tunnels (c). After that, the nylon thread was tied up (d). F, femur; T, tibia.

### Surgical Procedures

The DMM and CATR procedures were performed on the left knee joint using a combination anesthetic (medetomidine, 0.375 mg/kg; midazolam, 2.0 mg/kg; and butorphanol, 2.5 mg/kg). DMM surgery was performed on the left knee joint of the mice as previously described.^20^ CATR surgery was performed following the same procedure as DMM surgery (**Fig. 1B**) with the subsequent creation of bone tunnels in the distal femur and proximal tibia using a 25-gauge needle and insertion of 4-0 nylon threads threaded through the bone tunnels (**Fig. 1B, a,b**). The 4-0 nylon threads were tied and secured (**Fig. 1B, c,d**) to suppress the joint rotational instability in the DMM model. To reduce differences with intervention among the groups, a bone tunnel was also created in the DMM group, and a nylon thread was tied loosely to maintain joint instability.

### Tibial Rotational Test

We performed a tibial rotational test using a constant force spring (0.05 kgf; Sanko Spring Co., Ltd., Fukuoka, Japan) and a soft X-ray device (M-60; Softex Co., Ltd., Kanagawa, Japan) to assess the knee joint rotational instability. At 8 and 12 weeks after surgery, we evaluated the knee joints of mice with the intact femur, tibia, and foot. The femur and tibia were fixed with the knee joint at 90° flexion. In addition, we pierced the tibia with a 25-gauge needle to serve as a reference for changes in tibial rotation (**Fig. 2A**). The proximal tibia was pulled both laterally and medially with a 4-0 nylon thread. Radiographs were taken during the medial and lateral traction. We defined the change in angle between the needle and the vertical line of the soft x-ray device as rotational instability. Soft X-ray radiography was performed at 28 kV and 1.5 mA with an exposure time of 1 s. Digital images were acquired using a NAOMI digital X-ray sensor (RF Co. Ltd., Nagano, Japan). The change in the tibial rotation was quantified using a dedicated image analysis software (Image J; National Institutes of Health, Bethesda, MD, USA).

**Figure 2.**
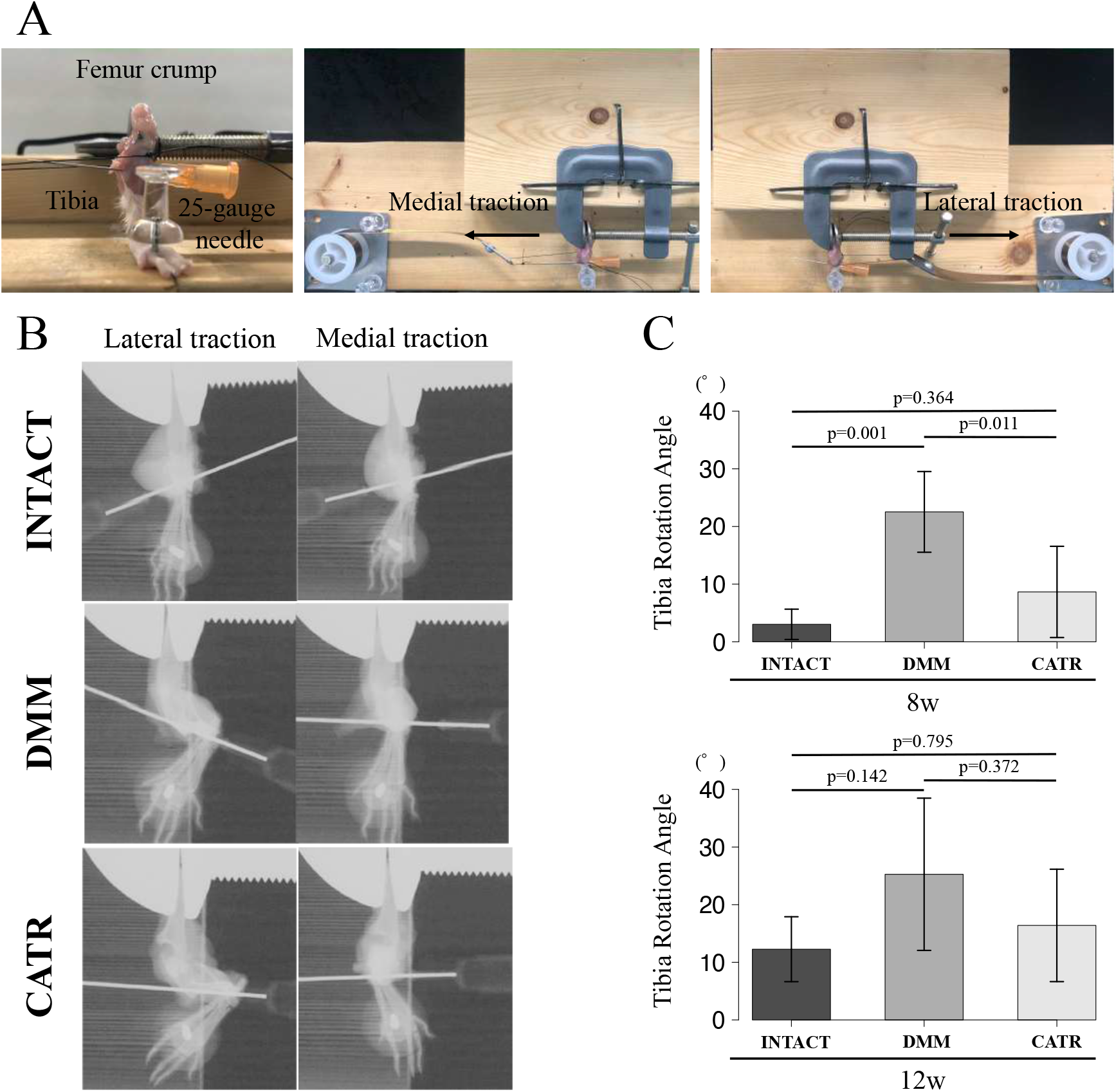
(A) System for tibial rotational test. The femur and tibia were set into the examination system at 90° flexion, and pierced the tibia with a 25-gauge needle. And rotational joint instability was maintained by medial and lateral traction. (B) Representative soft X-ray radiograph taken during the tibial rotational test on the knee joint. (C) Tibia rotation angle on the tibial rotational test. At 8 weeks, tibia rotation angle in the DMM group was significantly higher than that in the INTACT and CATR groups. At 12 weeks, DMM group was higher than that in the INTACT and CATR groups.

### Histological Analysis

Mice were sacrificed at 8 and 12 weeks after surgery, wherein 5 knees were assessed in each group for both time points. Subsequently, the knee joint was fixed in 4% paraformaldehyde/phosphate-buffered saline for 24 h, followed by decalcification in 10% ethylenediaminetetraacetic acid for 21 days, dehydration in 70% and 100% ethanol and xylene, and embedding in paraffin blocks. Thin sections (6 μm) were cut in the sagittal plane using a microtome (ROM-360; Yamato Kohki Industrial Co., Ltd., Saitama, Japan), stained with safranin-O/fast green, and subjected to histological evaluation to estimate the degree of cartilage and meniscal degeneration. The Osteoarthritis Research Society International (OARSI) histopathological grading system was used to assess cartilage degeneration for structural changes and fibrillation lesions.^23^ The mouse meniscus histological grading system established by Kwok et al.^21^ was used to evaluate the degeneration in the anterior horn of the meniscus. Two independent observers (KT and KM) performed OARSI scoring on an eight-stage scale (0, 0.5, 1-6) and meniscus scoring on a five-point scale (0-4), with the mean values retained.

### Immunohistochemical Analysis

To evaluate the expression of TNF-α and MMP-13, we performed immunohistochemical staining following the avidin-biotin complex method using the VECTASTAIN Elite ABC Rabbit IgG Kit (Vector Laboratories, Burlingame, CA, USA). The tissue sections were deparaffinized with xylene and ethanol, and antigen activation was performed using proteinase K (Worthington Biochemical Co., Lakewood, NJ, USA) for 15 min. Endogenous peroxidase was inactivated with 0.3% H_2_O_2_/methanol for 30 min. Nonspecific binding of the primary antibody was blocked using normal goat serum for 30 min, and then the sections were incubated with anti-TNF-α and anti-MMP-13 primary antibodies overnight at 4°C. Afterward, the sections were incubated with biotinylated secondary anti-rabbit IgG antibody and stained using Dako Liquid DAB+ Substrate Chromogen System (Dako, Glostrup, Denmark). Cell nuclei were stained with hematoxylin. For analysis, we calculated the ratio of the number of TNF-α- or MMP-13-positive cells to the number of chondrocytes in the articular cartilage area of 10 000 μm^2^ (100 μm × 100 μm).

### Statistical Analysis

All analyses were performed using R Studio version 1.2.5019. The normality of the data distribution was assessed using the Shapiro-Wilk test. Tibial rotation changes were normally distributed, while OARSI scores, meniscus scores, and percentages of TNF-α- and MMP-13-positive cells were not. Parametric data were compared using a one-way analysis of variance and post-hoc Tukey’s test. Non-parametric data were compared using the Kruskal-Wallis test and post-hoc Steel-Dwass analysis. All significance thresholds were set to 5%. Parametric data are presented as mean ± standard deviation, and nonparametric data are presented as median with interquartile ranges [IQR].

## RESULTS

### Evaluation of Joint Rotational Instability Using Soft X-ray Analysis

Joint rotational instability was quantified in terms of the change in tibial rotation angle, as determined using soft X-ray radiography with the tibial rotational test(**Fig. 2B**). At 8 weeks, the mean change in tibial rotation angle in the DMM group was significantly higher than that in the INTACT and CATR groups (INTACT,3.03 ± 2.63 degrees; DMM, 22.53 ± 7.00 degrees; CATR, 8.65 ± 7.91 degrees). At 12 weeks, although not significant, the mean change in tibial rotation angle in the DMM group was higher than that in the INTACT and CATR groups (INTACT, 12.27 ± 5.63 degrees; DMM, 25.27 ± 13.21 degrees; CATR, 16.40 ± 9.75 degrees). The exact P-values for each comparison are shown in **Figure 2C**.

### Histological Analysis

Representative histological images of the cartilage at 8 and 12 weeks are presented in **Figure 3A**. A total of 5 knees were assessed in each group for both time points. Structural changes in the surface layer of cartilage in the DMM group were observed after 8 and 12 weeks. At 12 weeks, mild degeneration of the cartilage surface layer was observed in the CATR group compared with that in the DMM group. In the meniscus, changes in the surface structure and decreased staining with safranin-O/fast green were observed in the DMM and CATR groups at 8 and 12 weeks (**Fig. 4A**).

**Figure 3.**
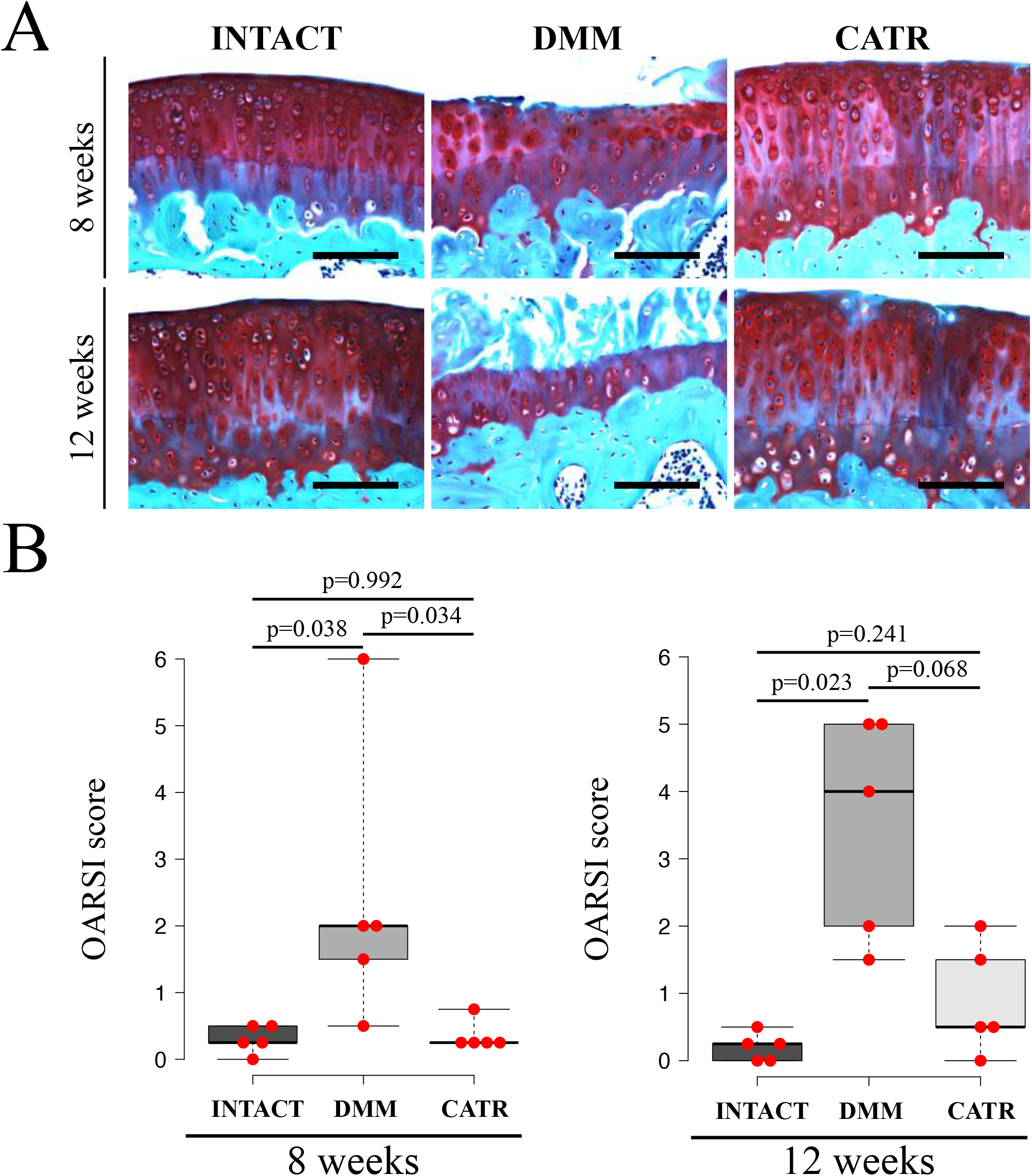
(A) Representative histological images of the cartilage at 8 and 12 weeks. (B) Results of the OARSI score. At 8 weeks, the OARSI scores significantly higher in the DMM group than in the INTACT and CATR groups. At 12 weeks, the OARSI scores in the DMM group tended to increase compared with those in the CATR group.

**Figure 4.**
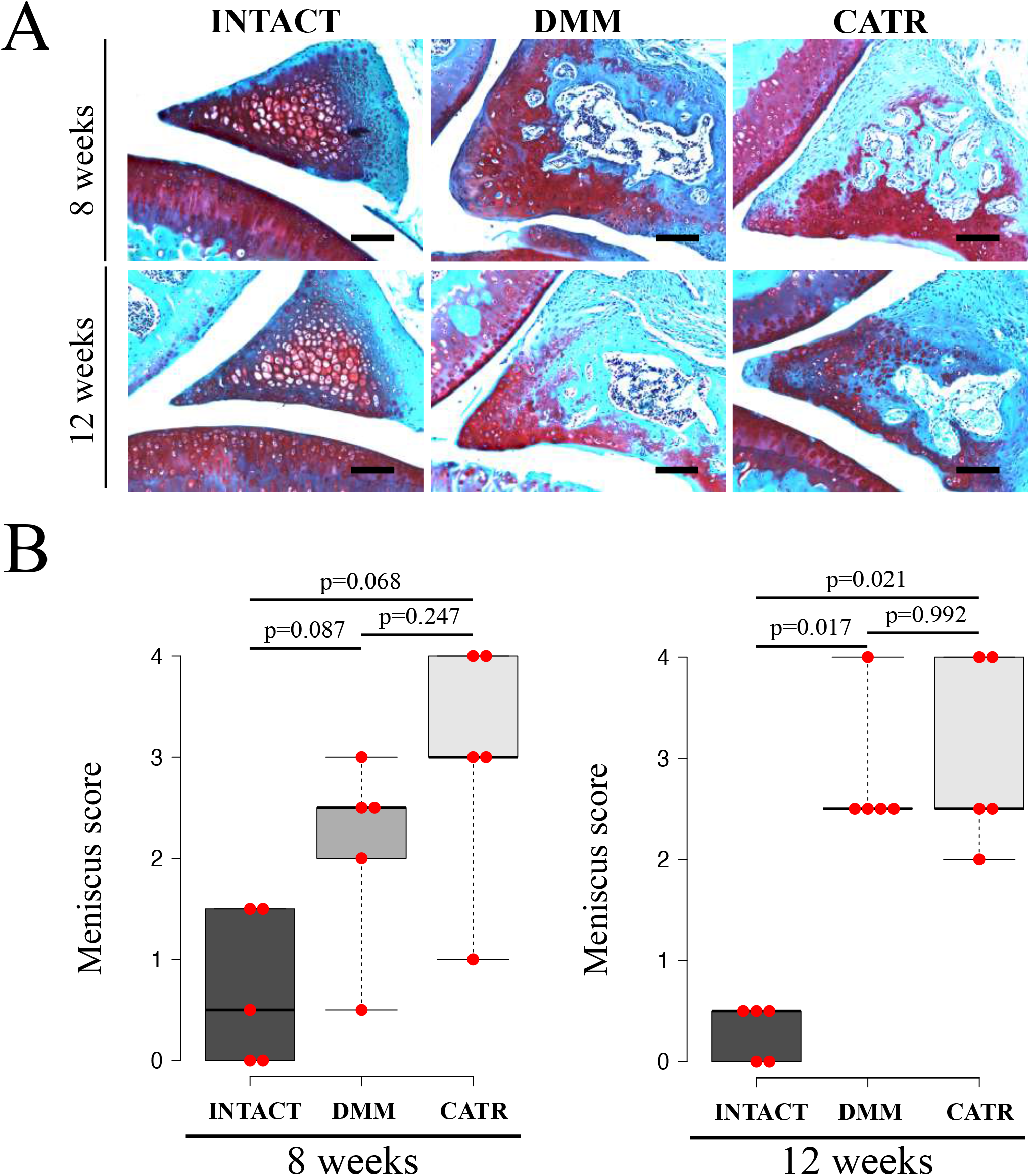
(A) Representative histological images of the meniscus at 8 and 12 weeks. (B) Results of the meniscus score. At 8 weeks, meniscal scores in the DMM and CATR groups tended to increase compared with those in the INTACT group. At 12 weeks, meniscal scores in the DMM and CATR groups were significantly higher than in the INTACT group.

Cartilage degeneration was assessed using the OARSI scores at 8 and 12 weeks (**Fig. 3B**). At 8 weeks, the OARSI scores significantly higher in the DMM group than in the INTACT and CATR groups (**Fig. 3B**) (DMM vs. INTACT, P = 0.038; DMM vs. CATR, P = 0.034), but there was no significant difference between the INTACT and CATR groups (**Fig. 3B**) (INTACT vs. CATR, P = 0.992). At 12 weeks, the OARSI scores in the DMM group were significantly higher than those observed in the INTACT group (**Fig. 3B**) (DMM vs. INTACT, P = 0.023). The OARSI scores in the DMM group tended to increase compared with those in the CATR group (**Fig. 3B**) (DMM vs. CATR, P = 0.068), although the difference was not significant. There was no significant difference between the INTACT and CATR groups (INTACT vs. CATR, P = 0.241) (**Fig. 3B**).

Evaluation of meniscal degeneration was performed using the mouse meniscus histological grading system established by Kwok et al.^21^ at 8 and 12 weeks (**Fig. 4B**). At 8 weeks, although not significant, meniscal degeneration scores in the DMM and CATR groups tended to increase compared with those in the INTACT group (**Fig. 4B**) (INTACT vs. DMM, P = 0.087; INTACT vs. CATR, P = 0.068). At 12 weeks, meniscal degeneration scores in the DMM and CATR groups were significantly higher than those observed in the INTACT group (INTACT vs. DMM, P = 0.017; INTACT vs. CATR, P = 0.021) (**Fig. 4B**). There was no significant difference between the DMM and CATR groups (DMM vs. CATR, P = 0.992) (**Fig. 4B**).

### Immunohistochemical Analysis

Representative images of TNF-α and MMP-13 immunostaining of cartilage specimens is shown in **Figure 5A**. At 8 weeks, the percentage of TNF-α-positive cells in the DMM group was significantly higher than that in the INTACT and CATR groups (INTACT vs. DMM, P = 0.024; DMM vs. CATR, P = 0.024) (**Fig. 5B**). Meanwhile, the percentage of MMP-13-positive cells in the DMM and CATR groups was significantly higher than in the INTACT group (INTACT vs. DMM, P = 0.024; INTACT vs. CATR, P = 0.024) (**Fig. 5B**). Although not significant, the percentage of MMP-13-positive cells in the DMM group tended to increase compared with that in the CATR group (DMM vs. CATR, P = 0.071). At 12 weeks, the percentage of TNF-α-positive cells in the DMM group was significantly higher than that in the INTACT and CATR groups (INTACT vs. DMM, P = 0.024; DMM vs. CATR, P = 0.024) (**Fig. 5B**). The percentage of TNF-α-positive cells in the CATR group was significantly higher than that in the INTACT group (CATR vs. INTACT, P = 0.043) (**Fig. 5B**). Similarly, the percentage of MMP-13-positive cells in the DMM group was significantly higher than that in the INTACT and CATR groups (INTACT vs. DMM, P = 0.024; DMM vs. CATR, P = 0.024) (**Fig. 5B**).

**Figure 5.**
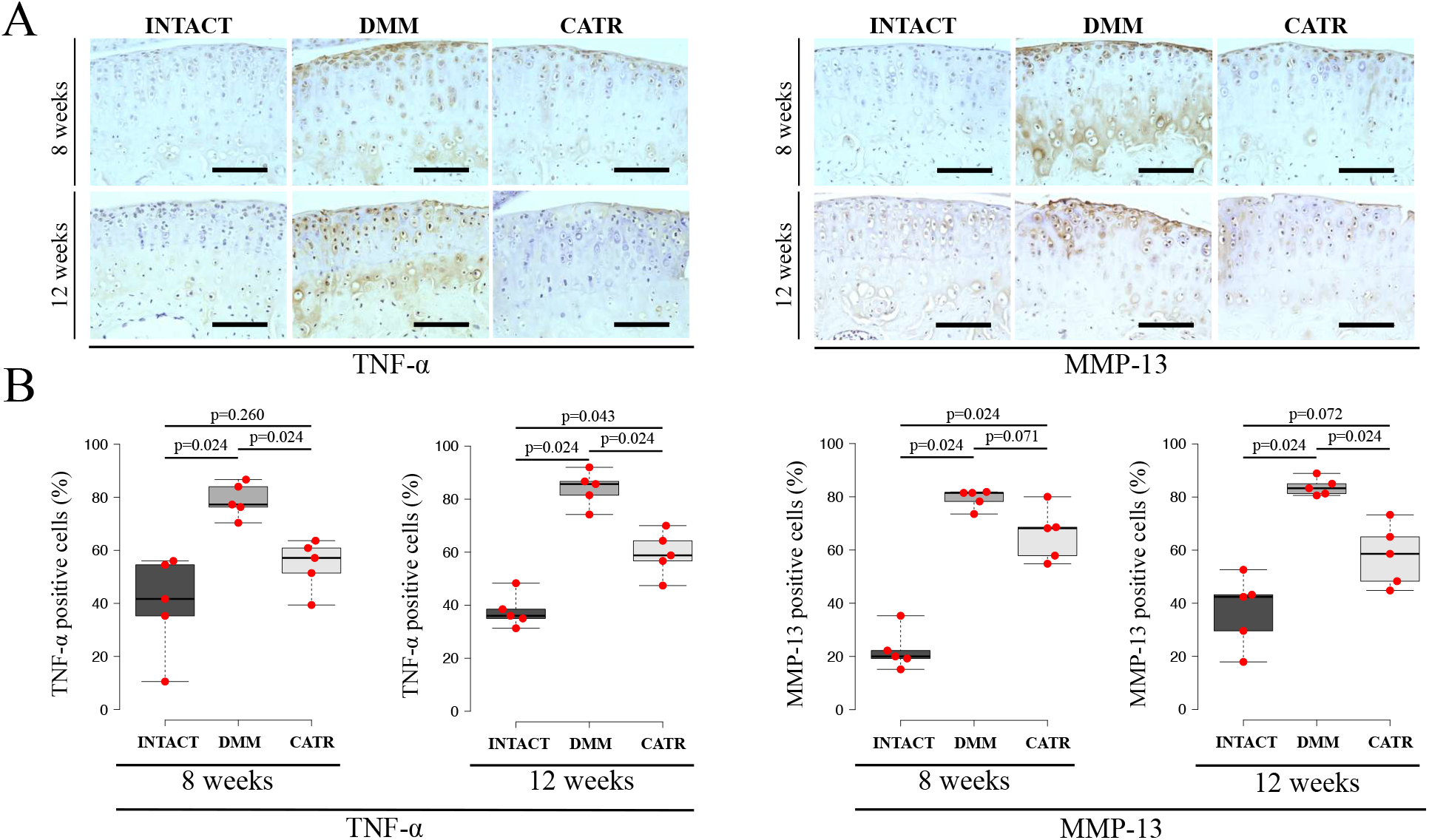
(A) Representative immunostaining images of TNF-α and MMP-13 at 8 and 12 weeks. (B) The percentage positive cell ratio of TNF-α and MMP-13. At 8 weeks, the percentage of TNF-α-positive cells in the DMM group was significantly higher than that in the INTACT and CATR groups. Although the percentage of MMP-13-positive cells in the DMM group tended to increase compared with that in the CATR group. At 12 weeks, the percentage of TNF-α-positive cells in the DMM group was significantly higher than that in the INTACT and CATR groups. And the percentage of MMP-13-positive cells in the DMM group was significantly higher than that in the INTACT and CATR groups.

## DISCUSSION

The data from this study showed that controlling the rotational instability caused by meniscal dysfunction suppresses articular cartilage degeneration, as confirmed by histological analysis. However, meniscal degeneration was not suppressed by controlling joint instability. Furthermore, the expression of inflammatory cytokines TNF-α and MMP-13 in articular cartilage was inhibited by controlling rotational instability, as confirmed by immunohistochemical analysis. These findings indicate that joint instability caused by meniscal dysfunction is a major contributory factor in the progression of knee OA.

In this study, we used the DMM model that causes dysfunction of the medial meniscus and the CATR model that controls the rotational instability that occurs in the DMM model. At 8 weeks, rotation instability occurred in the DMM model but was reduced in the CATR model, as evidenced by the results of the tibial rotation test using soft X-rays. At 12 weeks, results were similar to that at 8 weeks, but no significant difference was observed. The OARSI score, indicating the severity of articular cartilage degeneration, was lower in the CATR group, which showed suppressed joint instability, than in the DMM group, which showed rotational instability. Articular cartilage degeneration depends on the magnitude of joint instability.^10^ Furthermore, controlling the anterior-posterior joint instability that occurs in the rat anterior cruciate ligament-transection (ACL-T) model suppresses articular cartilage degeneration.^12,24^ Similarly, in this study, we found that the magnitude of joint instability affected the severity of articular cartilage degeneration. Our study findings suggest that in addition to the anterior-posterior joint instability that occurs in the ACL-T model, rotational instability is a factor that causes articular cartilage degeneration.

TNF-α expression promotes the catabolism of chondrocytes and disrupts chondrocyte homeostasis. TNF-α is an important inflammatory cytokine in OA and induces the production of several matrix metalloproteinases, such as MMP-13.^25–27^ MMP-13 induced by TNF-α degrades collagen type 2, the main extracellular matrix of articular cartilage.^28^ It has been reported that the expression of these cytokines that promote the catabolism of chondrocytes is reduced in the state of suppressed joint instability.^12,24^ In this study, the expression levels of TNF-α and MMP-13 in articular cartilage were lower in the CATR group than in the DMM group, suggesting that suppression of joint instability also inhibited the expression of TNF-α and MMP-13. As supported by our findings, interventions that control joint instability may be effective means of suppressing the expression of inflammatory cytokines and articular cartilage degeneration.

Notably, the expression of TNF-α and MMP-13 in the articular cartilage of the CATR group was higher than that in the INTACT group. Murata et al.^12^ and Onitsuka et al.^24^ reported that articular cartilage degeneration could be delayed by suppressing joint instability in the ACL-T model. However, the expression of TNF-α and MMP-13 also increased upon suppression of joint instability compared with that in the control group. Knee OA is a disease that involves not only mechanical stress such as joint instability but also various factors such as inflammation and immune response. In this study, the expression of TNF-α and MMP-13 in the articular cartilage region was not completely abolished. Although suppressing joint instability can reduce abnormal mechanical stress, further research is needed to elucidate its effects on other factors such as inflammation.

In our study, there was no significant difference in the meniscal degeneration scores between the DMM and CATR groups. At 12 weeks, the meniscal degeneration score was significantly higher in the DMM and CATR groups than in the INTACT group. Severe meniscal degeneration has been reported to already occur in the DMM model 2 weeks after surgical intervention.^21^ Approximately 70% of tissue weight in the meniscus is water, while the rest is made up of organic matter, mainly extracellular matrix and cells. Collagen makes up the majority of organic matter, followed by glycosaminoglycans.^29,30^

The inner region of the meniscus has a higher percentage of collagen type 2 than the outer region, which has a composition similar to those of the articular cartilage.^31^ With these biochemical properties, it has been shown that the semilunar plates play a role in load-bearing, load transfer, and shock absorption, as well as resisting axial compressive loading.^32,33^ In the DMM and CATR groups, we observed a decreased staining of safranin-O/fast green with concomitant structural changes in the meniscus at 8 and 12 weeks after surgery. Therefore, it is inferred that the load distribution in these two groups was not optimal, and the compressive load increased.

Further, meniscal degeneration was observed in both the DMM and CATR groups, but articular cartilage degeneration was more severe in the DMM group. These results suggest that articular cartilage degeneration in the DMM group is not caused by load distribution disruption and compressive load increase due to meniscal dysfunction but is related to joint instability. Our findings support our hypothesis that suppression of joint instability is a common and effective intervention target in articular cartilage degeneration.

This study has significant limitations. Joint instability was the focus of our study. We showed that the CATR model suppresses the rotational instability that occurs in the DMM model. However, we only verified static instability and not dynamic instability. Thus, it is necessary to further confirm whether there is a difference in joint instability between the DMM and CATR models in a dynamic environment, such as walking. In addition, OA is considered a disease of the entire joint, characterized by articular cartilage degeneration, synovitis, and changes in the subchondral bone. In particular, subchondral bone and cartilage have been found to have biological interactions.^34–36^ In this study, the effect of joint instability on articular cartilage degeneration was clarified, but its effect on the subchondral bone is unclear. Further research requires analysis of surrounding tissues.

In conclusion, articular cartilage degeneration was effectively suppressed by controlling the rotational instability caused by meniscal dysfunction. Our findings suggest that control of joint instability is a good strategy for minimizing articular cartilage degeneration and that joint instability from meniscal dysfunction is likely a major contributory factor to the articular cartilage degeneration in the DMM model. Meniscal degeneration and dysfunction are involved in the progression of OA.^17–19^ Thus, our study suggests that suppression of rotational instability in the knee joint is an effective therapeutic measure for preventing OA progression.

## Author Contributions

All authors approved the final version to be published.

Study design: KA, KM, NK, and TK

Data collection, Histological analysis: KA, KT, YO, KO, K Morosawa and SE

Manuscript composition KA, KM, NK, TK

## ACKNOWLEDGEMENTS

The author(s) received no financial support for the research, authorship, and/or publication of this article.

## DECLARATION OF CONFLICTING INTERESTS

The authors declare that there is no conflict of interest

## Ethical Approval

This study was approved by the Animal Research Committee of Saitama Prefectural University (Approval Number: 29-13).

## Animal Welfare

This study followed institutional guidelines for humane animal treatment and complied with relevant legislation.

## Notes

### Competing Interest Statement

The authors have declared no competing interest.

